## Supplementary R Code for "Chemical patterns of colony membership and mother-offspring similarity in Antarctic fur seals are reproducible over time"

*April 2020*

### Used packages

Install with "install.packages". After installation, packages can be called with 'library' oder 'require'

```
library(GCalignR)
library(vegan)
library(readr)
library(ggplot2)
library(ggbeeswarm)
library(tidyverse)
library(pairwiseAdonis)
```

### Alignment and preliminary data properties

```
## Load and view GCalignR alignment objects for GCMS scent data
## in two and six breeding beaches
load("RData/objects/mom_pup_alignment_GCalignR.RData")
mom_pup_aligned
```

```
## Summary of Peak Alignment running align_chromatograms
## Input: all_dfs[index2]
## Start:  2019-05-21 11:41:08  Finished:  2019-05-21 11:44:23
##
## Call:
##   GCalignR::align_chromatograms(data=[, data=all_dfs, data=index2, rt_col_name=RT,
##   rt_cutoff_low=15, rt_cutoff_high=54.7, reference=P13, max_linear_shift=0.05,
##   max_diff_peak2mean=0.08, min_diff_peak2peak=0.03, delete_single_peak=T,
##   sep=\t, ...=)
##
## Summary of scored substances:
##    total singular retained
##      157       39      118
##
## In total 157 substances were identified among all samples. 39 substances were
##   present in just one single sample and were removed. 118 substances are retained
##   after all filtering steps.
##
## Sample overview:
##   The following 101 samples were aligned to the reference 'P13':
##   M01, M02, M03, M04, M05, M06, M07, M08, M09, M10, M11, M12, M13, M14, M15, M16,
##   M17, M18, M19, M20, M21, M22, M23, M24, M25, M26, M27, M28, M29, M30, M31, M32,
##   M33, M34, M35, M36, M37, M38, M39, M40, M41, M42, M43, M44, M45, M46, M47, M48,
```

```
##   M49, M50, P01, P02, P03, P04, P05, P06, P07, P07b, P08, P09, P10, P11, P12, P13,
##   P14, P15, P16, P17, P18, P19, P20, P21, P22, P23, P24, P25, P26, P27, P28, P29,
##   P30, P31, P32, P33, P34, P35, P36, P37, P38, P39, P40, P41, P42, P43, P44, P45,
##   P46, P47, P48, P49, P50
##
## For further details type:
##   'gc_heatmap(x)' to retrieve heatmaps
##   'plot(x)' to retrieve further diagnostic plots
```

```r
load("RData/objects/pup_colonies_alignment_GCalignR.RData")
pup_colonies_aligned
```

```
## Summary of Peak Alignment running align_chromatograms
## Input: all_dfs[index4]
## Start:  2019-05-23 11:36:39  Finished:  2019-05-23 11:40:15
##
## Call:
##   GCalignR::align_chromatograms(data=[, data=all_dfs, data=index4, rt_col_name=RT,
##   rt_cutoff_low=15, rt_cutoff_high=54.7, reference=P13, max_linear_shift=0.05,
##   max_diff_peak2mean=0.08, min_diff_peak2peak=0.03, delete_single_peak=T,
##   sep=\t, ...=)
##
## Summary of scored substances:
##    total singular retained
##      143       28      115
##
## In total 143 substances were identified among all samples. 28 substances were
##   present in just one single sample and were removed. 115 substances are retained
##   after all filtering steps.
##
## Sample overview:
##   The following 110 samples were aligned to the reference 'P13':
##   P01, P02, P03, P04, P05, P06, P07, P07b, P08, P09, P10, P100, P101, P102, P103,
##   P104, P105, P106, P107, P108, P109, P11, P12, P13, P14, P15, P16, P17, P18, P19,
##   P20, P21, P22, P23, P24, P25, P26, P27, P28, P29, P30, P31, P32, P33, P34, P35,
##   P36, P37, P38, P39, P40, P41, P42, P43, P44, P45, P46, P47, P48, P49, P50, P51,
##   P52, P53, P54, P55, P56, P57, P58, P59, P60, P61, P62, P63, P64, P65, P66, P67,
##   P68, P69, P70, P71, P72, P73, P74, P75, P76, P77, P78, P79, P80, P81, P82, P83,
##   P84, P85, P86, P87, P88, P89, P90, P91, P92, P93, P94, P95, P96, P97, P98, P99
##
## For further details type:
##   'gc_heatmap(x)' to retrieve heatmaps
##   'plot(x)' to retrieve further diagnostic plots
```

```r
## Load raw information for all samples containing
## raw peaks and calculate mean peak number
load("RData/objects/seal_raw_dfs.Rdata")

individual_peak_number <- NULL
for (i in 1:length(seal_dfs.list)) {
  individual_peak_number[i] <- length(seal_dfs.list[[i]]$RT)
}

mean_ind_peaks <- mean(individual_peak_number)
sd_ind_peaks <- sd(individual_peak_number)
```

```r
cat("\n", "\n", "Mean peaks:", as.character(mean_ind_peaks), "\n", "Peak SD:",
    as.character(sd_ind_peaks))
```

```
##
##
##  Mean peaks: 34.175
##  Peak SD: 10.8445563540296
```

#### NMDS scaling of mother-pup alignment data

```r
load("RData/objects/mom_pup_alignment_GCalignR.RData")

scent_factors_raw <- read_delim("documents/metadata_seal_scent.txt",
                                "\t", escape_double = FALSE, trim_ws = TRUE)
scent_factors_raw <- as.data.frame(scent_factors_raw[-c(194:209),])

# set sample names as row names, ensure there are no duplicates
scent_factors <- scent_factors_raw[,-1]
rownames(scent_factors) <- scent_factors_raw[,1]

## check for empty samples, i.e. no peaks
x <- apply(mom_pup_aligned$aligned$RT, 2, sum)
x <- which(x == 0)

## normalise area and return a data frame
scent <- norm_peaks(mom_pup_aligned, conc_col_name = "Area",rt_col_name = "RT",
                    out = "data.frame")
## common transformation for abundance data to reduce the extent of mean-variance trends
scent <- log(scent + 1)

## subset scent_factors
scent_factors <- scent_factors[rownames(scent_factors) %in% rownames(scent),]
scent <- scent[rownames(scent) %in% rownames(scent_factors),]

## keep order of rows consistent
scent <- scent[match(rownames(scent_factors),rownames(scent)),]

## get number of compounds for each individual sample after alignment
num_comp <- as.vector(apply(scent, 1, function(x) length(x[x>0])))

## bray-curtis similarity
scent_nmds.obj <- vegan::metaMDS(comm = scent, k = 2, try = 999,
                                 trymax = 9999, distance = "bray")

scent_nmds <- as.data.frame(scent_nmds.obj[["points"]])

scent_nmds <- cbind(scent_nmds,
                    age = scent_factors[["age"]],
                    tissue_tag = scent_factors[["tissue_tag"]],
                    colony = scent_factors[["colony"]],
                    family = as.factor(scent_factors[["family"]]),
                    clr = as.factor(scent_factors[["clr"]]),
                    shp = as.factor(scent_factors[["shp"]]),
```

```
                     gcms = as.factor(scent_factors[["gcms_run"]]),
                     peak_res = as.factor(scent_factors[["peak_res"]]),
                     sample_qlty = as.factor(scent_factors[["sample_qlty"]]),
                     vialdate = as.factor(scent_factors[["gcms_vialdate"]]),
                     captured = as.factor(scent_factors[["capture_date"]]),
                     sex = scent_factors[["sex"]],
                     num_comp = num_comp)
scent_nmds <- scent_nmds %>% mutate(BeachAge = str_c(colony, age, sep = "_"))
# creates & adds new variable BeachAge and simplifies plotting
```

#### Colony and family membership in SSB and FWB mom-pup pairs

```
load("RData/objects/mom_pup_nmds_scaling.RData")

## colony membership plot
mp_colony_gg <- ggplot(data = scent_nmds) +
  geom_point(size = 4.5, aes(MDS1, MDS2, color = BeachAge, shape = BeachAge)) +
  scale_shape_manual(values = c(19, 1, 19, 1),
                     labels = c("FWB mothers ", "FWB pups ",
                                "SSB mothers ", "SSB pups ")) +
  scale_color_manual(values = c("#D55E00", "#D55E00", "#56B4E9", "#56B4E9"),
                     labels = c("FWB mothers ", "FWB pups ",
                                "SSB mothers ", "SSB pups ")) +
  theme_void() +
  ylim(-0.75,1.1) +
  annotate("text", x = 0.64, y = 1.1, label = "A", size = 5) +
  annotate("text", x = 0.47, y = -0.74, label = "2D Stress: 0.23", size = 4) +
  theme(panel.background = element_rect(colour = "black", size = 1, fill = NA),
        aspect.ratio = 1,
        legend.position = "none",
        legend.title = element_blank(),
        legend.background = element_rect(size = 0.3, linetype = "solid", color = "black"))
# call colony membership plot
mp_colony_gg
```

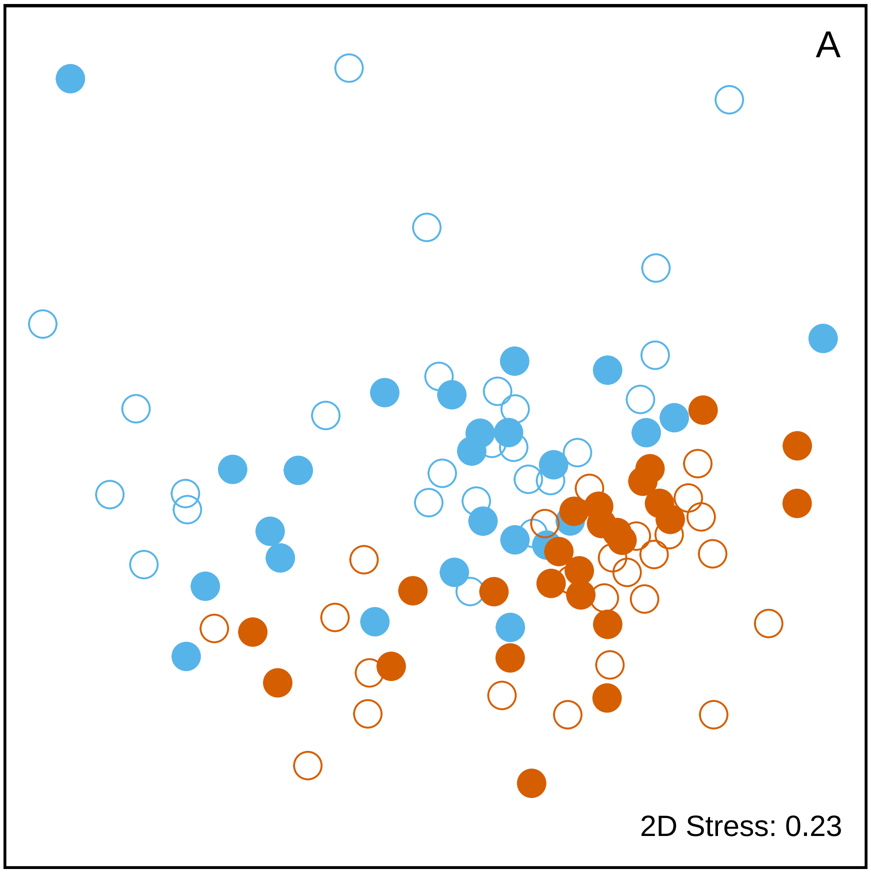


```r
## mother-offspring similarity plot
# create color palette for the plot
clr <- c("#D55E00", "red", "#56B4E9", "#009E73","#000000", "#CC79A7")

# assign pch values for plotting
shp <- c(0,1,2,7,10,5,6,18,16,17,15)

# create unique color-pch pairs
color_shape_pairs <- crossing(clr,shp)

# randomly sample 50 unique pairs (sample without replacement)
set.seed(123) # always get same pairs in a run
color_shape_pairs <- color_shape_pairs[sample(nrow(color_shape_pairs), 50),]

# assign new dataframes to transform scent_nmds$clr & shp with the unique values we created
color_shape_pairs_plot <- rbind(color_shape_pairs[1:25,],color_shape_pairs[1:7,]
                                ,color_shape_pairs[7,],  color_shape_pairs[8:25,],
                                color_shape_pairs[26:50,], color_shape_pairs[26:50,])
scent_nmds$clr <- as.factor(color_shape_pairs_plot$clr)
scent_nmds$shp <- as.factor(color_shape_pairs_plot$shp)

# call family plot
mp_family_gg <- ggplot(data = scent_nmds,aes(MDS1,MDS2, color = clr, shape = shp)) +
  geom_point(size = 4.5) +
  scale_shape_manual(values = as.numeric(levels(scent_nmds$shp))) +
  theme_void() +
```

```
  ylim(-0.75,1.1) +
  scale_color_manual(values = levels(scent_nmds$clr)) +
  annotate("text", x = 0.64, y = 1.1, label = "B", size = 5) +
  annotate("text", x = 0.48, y = -0.74, label = "2D Stress: 0.23", size = 4) +
  theme(panel.background = element_rect(colour = "black", size = 1,
                                      fill = NA), aspect.ratio = 1,
        legend.position = "none")
mp_family_gg
```

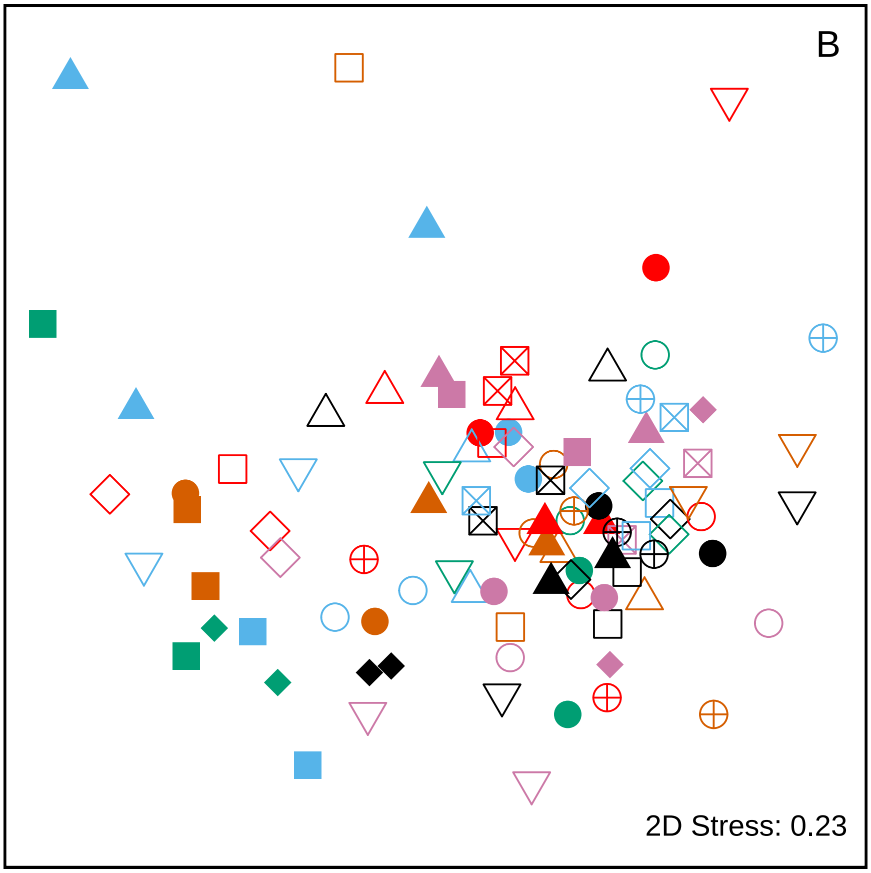


#### PERMANOVA for mother-pup similiarity and colony membership

```
# set seed to reproduce the same outcome (can vary due to different permutations!)
set.seed(123)
adonis(scent ~ age+colony+colony:family,
       data = scent_factors,
       method = "bray",
       permutations = 99999)
```

```
##
## Call:
## adonis(formula = scent ~ age + colony + colony:family, data = scent_factors,      permutations = 9999
##
## Permutation: free
## Number of permutations: 99999
##
```

```
## Terms added sequentially (first to last)
##
##                Df SumsOfSqs MeanSqs F.Model      R2  Pr(>F)
## age             1    0.3217 0.32170  2.6896 0.02253 0.00421 **
## colony          1    1.0847 1.08475  9.0692 0.07599   1e-05 ***
## colony:family   2    1.3870 0.69351  5.7982 0.09716   1e-05 ***
## Residuals      96   11.4823 0.11961         0.80432
## Total         100   14.2758                 1.00000
## ---
## Signif. codes:  0 '***' 0.001 '**' 0.01 '*' 0.05 '.' 0.1 ' ' 1
```

```
# test for group dispersal
mod <- betadisper(vegdist(scent), scent_factors$colony, type = "median")
anova(mod)
```

```
## Analysis of Variance Table
##
## Response: Distances
##           Df Sum Sq  Mean Sq F value  Pr(>F)
## Groups     1 0.0442 0.044201   5.136 0.02561 *
## Residuals 99 0.8520 0.008606
## ---
## Signif. codes:  0 '***' 0.001 '**' 0.01 '*' 0.05 '.' 0.1 ' ' 1
```

#### NMDS scaling and colony membership in six pup colonies

```
load("RData/objects/pup_colonies_alignment_GCalignR.RData")
scent_factors_raw <- read_delim("documents/metadata_seal_scent.txt",
                                "\t", escape_double = FALSE, trim_ws = TRUE)
scent_factors_raw <- as.data.frame(scent_factors_raw[-c(194:209),])

# set sample names as row names, ensure there are no duplicates
scent_factors <- scent_factors_raw[,-1]
rownames(scent_factors) <- scent_factors_raw[,1]

## check for empty samples, i.e. no peaks
x <- apply(pup_colonies_aligned$aligned$RT, 2, sum)
x <- which(x == 0)

## normalise area and return a data frame
scent <- norm_peaks(pup_colonies_aligned, conc_col_name = "Area",rt_col_name = "RT",
                    out = "data.frame")
## common transformation for abundance data to reduce the extent of mean-variance trends
scent <- log(scent + 1)

## subset scent_factors
scent_factors <- scent_factors[rownames(scent_factors) %in% rownames(scent),]
scent <- scent[rownames(scent) %in% rownames(scent_factors),]

## keep order of rows consistent
scent <- scent[match(rownames(scent_factors),rownames(scent)),]

## get number of compounds for each individual sample after alignment
num_comp <- as.vector(apply(scent, 1, function(x) length(x[x>0])))
```

```
## bray-curtis similarity
scent_nmds.obj <- metaMDS(comm = scent, k = 2, try = 999,
                          trymax = 9999, distance = "bray")
## MDS outcome evaluated with PCA for factor colony in metadata table for individuals
scent_nmds <- with(scent_factors, MDSrotate(scent_nmds.obj, colony))

## get x and y coordinates
scent_nmds <- as.data.frame(scent_nmds[["points"]])

## add the colony as a factor to each sample
scent_nmds <- cbind(scent_nmds,
                    age = scent_factors[["age"]],
                    tissue_tag = scent_factors[["tissue_tag"]],
                    colony = scent_factors[["colony"]],
                    family = as.factor(scent_factors[["family"]]),
                    clr = as.factor(scent_factors[["clr"]]),
                    shp = as.factor(scent_factors[["shp"]]),
                    gcms = as.factor(scent_factors[["gcms_run"]]),
                    peak_res = as.factor(scent_factors[["peak_res"]]),
                    sample_qlty = as.factor(scent_factors[["sample_qlty"]]),
                    vialdate = as.factor(scent_factors[["gcms_vialdate"]]),
                    captured = as.factor(scent_factors[["capture_date"]]),
                    sex = scent_factors[["sex"]],
                    num_comp = num_comp)
# creates & adds new variable BeachAge
scent_nmds <- scent_nmds %>% mutate(BeachAge = str_c(colony, age, sep = "_"))
```

Colony membership plot for six pup colonies (Supplementary figure)

```
load("RData/objects/pup_colonies_nmds_scaling.RData")

pup_colony_gg <- ggplot(data = scent_nmds, aes(MDS1, MDS2, color = colony, shape = colony)) +
  geom_point(size = 4.5) +
  scale_shape_manual(values = c(15,20,17,15,17,18),
                     labels = c("FWB", "Johnson cove", "Landing beach", "Main bay", "Natural arch", "SS
  scale_color_manual(values = c("#D55E00", "#000000", "#E69F00", "#009E73", "#CC79A7", "#0072B2"),
                     labels = c("FWB", "Johnson cove", "Landing beach", "Main bay", "Natural arch", "SS
  theme_void() +
  annotate("text", x = 0.6, y = -0.94, label = "2D Stress: 0.24", size = 4) +
  theme(panel.background = element_rect(colour = "black", size = 1, fill = NA),
        aspect.ratio = 1,
        legend.position = "right", #c(0.1,0.87),
        legend.title = element_blank(),
        # legend.key.size = unit(0.5, "cm"),
        # legend.key.width = unit(0.5, "cm"),
        legend.background = element_rect(size = 0.3, linetype = "solid", color = "black"))

pup_colony_gg
```

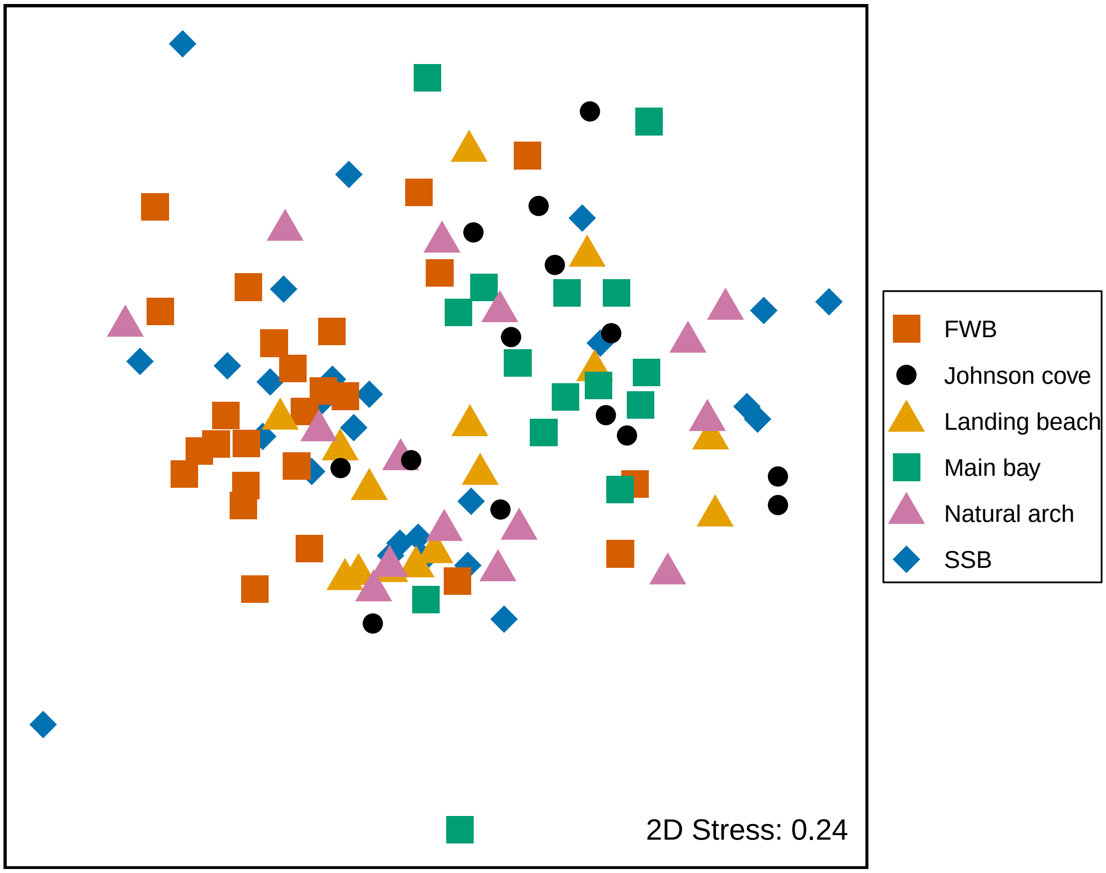


PERMANOVA for colony membership in six pup colonies

```
set.seed(123)
adonis(scent ~ age+colony+colony:family,
       data = scent_factors,
       permutations = 99999)
```

```
##
## Call:
## adonis(formula = scent ~ age + colony + colony:family, data = scent_factors,      permutations = 9999
##
## Permutation: free
## Number of permutations: 99999
##
## Terms added sequentially (first to last)
##
##                 Df SumsOfSqs MeanSqs F.Model      R2 Pr(>F)
## colony           5    3.1874 0.63749  5.1748 0.19128  1e-05 ***
## colony:family    6    1.4037 0.23395  1.8991 0.08424  7e-05 ***
## Residuals       98   12.0727 0.12319         0.72449
## Total          109   16.6639                 1.00000
## ---
## Signif. codes:  0 '***' 0.001 '**' 0.01 '*' 0.05 '.' 0.1 ' ' 1
```

```
# pairwise PERMANOVA
pairwiseAdonis::pairwise.adonis(scent, scent_factors$colony, perm = 99999)
```

```
##                              pairs Df SumsOfSqs  F.Model         R2 p.value
```

```
## 1                     SSB vs FWB  1 0.8086332 6.080018 0.11038519 0.00001
## 2           SSB vs landing_beach  1 0.4880064 3.544865 0.08332062 0.00081
## 3                SSB vs main_bay  1 0.8584284 6.181387 0.13681269 0.00001
## 4            SSB vs natural_arch  1 0.5667911 4.172853 0.09665456 0.00010
## 5                 SSB vs johnson  1 0.6168108 4.407128 0.10392422 0.00002
## 6           FWB vs landing_beach  1 0.5117674 4.168468 0.09885273 0.00006
## 7                FWB vs main_bay  1 0.9039828 7.289582 0.16095494 0.00001
## 8            FWB vs natural_arch  1 0.9556981 7.905832 0.17221847 0.00001
## 9                 FWB vs johnson  1 0.8911846 7.145352 0.16185966 0.00003
## 10     landing_beach vs main_bay  1 0.4229462 3.322412 0.10607138 0.00201
## 11 landing_beach vs natural_arch  1 0.3694950 3.002564 0.09684889 0.00421
## 12      landing_beach vs johnson  1 0.3459312 2.694198 0.09073145 0.01179
## 13      main_bay vs natural_arch  1 0.7381617 5.917530 0.17446818 0.00001
## 14           main_bay vs johnson  1 0.4083747 3.137902 0.10411813 0.00034
## 15       natural_arch vs johnson  1 0.3053377 2.428242 0.08251399 0.01630
##    p.adjusted sig
## 1     0.00015  **
## 2     0.01215   .
## 3     0.00015  **
## 4     0.00150   *
## 5     0.00030  **
## 6     0.00090  **
## 7     0.00015  **
## 8     0.00015  **
## 9     0.00045  **
## 10    0.03015   .
## 11    0.06315
## 12    0.17685
## 13    0.00015  **
## 14    0.00510   *
## 15    0.24450
```

```
# test for group dispersal
mod2 <- betadisper(vegdist(scent), scent_factors$colony, type = "median")
anova(mod2)
```

```
## Analysis of Variance Table
##
## Response: Distances
##            Df  Sum Sq   Mean Sq F value Pr(>F)
## Groups      5 0.02003 0.0040065   0.497 0.7779
## Residuals 104 0.83841 0.0080616
```

### Re-evaluation of 2011 field season scent data

Perform non-metric multidimensional scaling

Re-evalution in PERMANOVA instead of ANOSIM

```
## PERMANOVA
set.seed(123)
adonis(scent ~ age+colony+colony:family,
       data = peak_factors,
       permutations = 99999)
```

```
##
```

```
## Call:
## adonis(formula = scent ~ age + colony + colony:family, data = peak_factors,      permutations = 9999
##
## Permutation: free
## Number of permutations: 99999
##
## Terms added sequentially (first to last)
##
##                Df SumsOfSqs MeanSqs F.Model      R2 Pr(>F)
## age             1    0.2014 0.20143  0.9785 0.01013 0.4613
## colony          1    2.5430 2.54300 12.3538 0.12790  1e-05 ***
## colony:family  2    1.2880 0.64400  3.1285 0.06478  1e-05 ***
## Residuals      77   15.8503 0.20585         0.79719
## Total          81   19.8827                 1.00000
## ---
## Signif. codes:  0 '***' 0.001 '**' 0.01 '*' 0.05 '.' 0.1 ' ' 1
```

```
# Test for heterogeneity
anova(betadisper(vegdist(scent), peak_factors$colony))
```

```
## Analysis of Variance Table
##
## Response: Distances
##           Df   Sum Sq   Mean Sq F value Pr(>F)
## Groups     1 0.000791 0.0007913   0.222 0.6388
## Residuals 80 0.285197 0.0035650
```

#### Effect size estimate by PERMANOVA $R^2$ bootstrap

```
## Load data and assign data to data.frames
load("RData/objects/R2_initial_season_btrap.RData")

old_season_colony <- paov_r2_results[[1]][[2]]
old_season_family <- paov_r2_results[[1]][[3]]

load("RData/objects/R2_replication_season_btrap.RData")
new_season_colony <- paov_r2_results[[1]][[2]]
new_season_family <- paov_r2_results[[1]][[3]]

MP_effectsize <- c(old_season_colony, new_season_colony,
                   old_season_family, new_season_family)

MP_effectsize.groups <- c(rep("Colony S1", 5000),
                          rep("Colony S2", 5000),
                          rep("Family S1", 5000),
                          rep("Family S2", 5000))

MP_effectsize.df <- data.frame(btrap_combined_results = MP_effectsize,
                               btrap_subset_groups = MP_effectsize.groups)
```

Effect size estimate plot

```
load("RData/objects/effect_size_df.RData")
# point estimates for PERMANOVA on non-bootstrapped (original) data
point_estimate <- c(0.1444734, 0.09168289, 0.08780086, 0.1209394)
```

```
# point estimate groups for reasons of comprehensibility
point_estimate_groups <- c("Colony S1", "Colony S2", "Family S1", "Family S2")

# plot commands
MP_effectsize_gg <- ggplot(MP_effectsize.df, aes(y = btrap_combined_results,
                                                  x = btrap_subset_groups,
                                                  color = btrap_subset_groups)) +
  # this arranges the points according to their density
  geom_quasirandom(alpha = 0.06, size = 3, width = 0.3, bandwidth = 1) +
  scale_color_manual(values = c("#E69F00" ,"#E69F00" ,"#CC79A7", "#CC79A7")) +
  # makes the boxplots
  geom_boxplot(width = 0.35, outlier.shape = NA, color = "white", alpha = 0.1, lwd=0.8) +
  annotate("point", x = 1, y = point_estimate[4], colour = "#000000",
           fill = "#CCCCCC", size = 2, shape = 21) +
  annotate("point", x = 2, y = point_estimate[3], colour = "#000000",
           fill = "#CCCCCC", size = 2, shape = 21) +
  annotate("point", x = 3, y = point_estimate[2], colour = "#000000",
           fill = "#CCCCCC", size = 2, shape = 21) +
  annotate("point", x = 4, y = point_estimate[1], colour = "#000000",
           fill = "#CCCCCC", size = 2, shape = 21) +
  # this is a possible theme of the plot, there are many others
  theme_classic() +
  # changes the labels on the x axis
  scale_y_continuous(limits = c(-0.01 ,0.25),
                     breaks = seq(0, 0.25, 0.05)) +
  scale_x_discrete(labels = c("Family S2" = "Mother-offspring similarity\nreplication study",
                              "Family S1" = "Mother-offspring similarity\noriginal study",
                              "Colony S2" = "Colony membership\nreplication study",
                              "Colony S1" = "Colony membership\noriginal study"),
                   limits = c("Family S2",
                              "Family S1",
                              "Colony S2",
                              "Colony S1")) +
  # geom_hline(yintercept = 0, linetype = "dashed") +
  xlab("") +
  # label for y axis
  ylab("Explained variation [R²]") +
  # flips plot so everything is horizontal
  coord_flip() +
  # adjust theme specifics
  theme(panel.background = element_rect(colour = "black", size = 1.25, fill = NA),
        text = element_text(size = 15),
        axis.text = element_text(colour = "black"),
        legend.position = "none")


MP_effectsize_gg
```

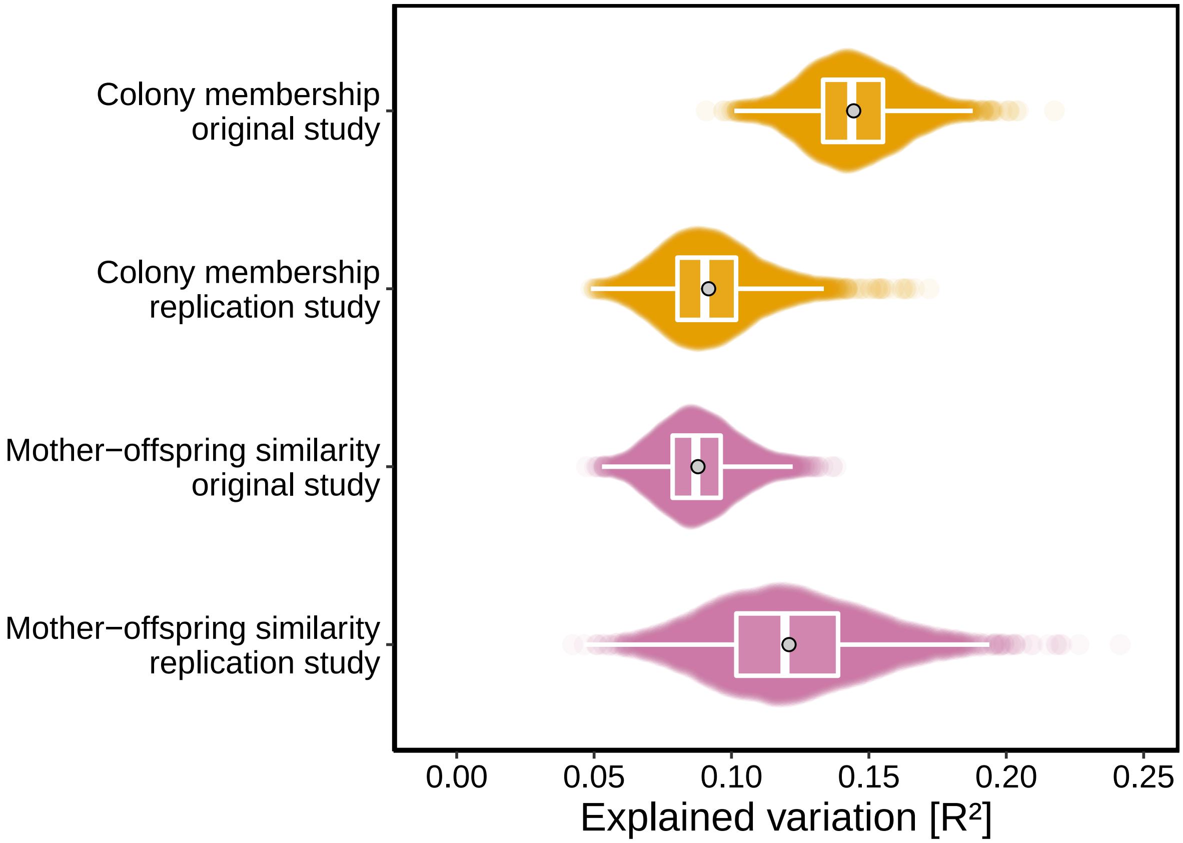


#### R2 Bootstrap Code

```r
## creates function 'scent_btrap_r2_swarm_data' that performs bootstrap

# Bootstrap to track R2 values for randomized subsets. In addition,
# bootstrap cannot only be used to randomize the chemical data frame
# to evaluate R2 distribution as effect size estimates,
# but also to evaluate R2 change for different subsets based on different
# premises. 1) Frequent peaks 2) Strong concentrations 3) Peaks identified by SIMPER

require(vegan)

# path: file path to scent_nmds-mompup2017_ssbfwb.RData",
#objects: scent_nmds, scent_nmds.obj, scent_factors, scent
# df.permutations: number of times the scent.df from loaded data will be permuted
# nmds.permutations: number of permutation in nMDS using Bray-Curtis
# btrap.iterations: number of procedure repeats

scent_btrap_r2_swarm_data <- function(path, df.permutations = 15,
                                      nmds.permutations = 999,
                                      btrap.iterations = 5000){
  # Create a data frame by permuting the data for scent
  # compounds data and also ensure that each population*age occur
  # same amounts of time in the permutation data frame.
  #----------------------------------------
```

```r
# load data frame with data of aligned fur seal chromatograms
load(path)
scent_factors <- peak_factors
# transfer BeachAge Column from scent_nmds to meta data.frame scent_factors
scent_factors <- cbind(scent_factors,
                       BeachAge = scent_nmds$BeachAge)

# create index column for meta data frame
scent_factors <- cbind(scent_factors,
                       SampleIndex = 1:length(rownames(scent_factors)))


# create data.frame to track PERMANOVA results over repeated tests
nonsubset_results_paov <- data.frame(R2_age = double(), p_colfam = double(),
                                     R2_residual = double(),
                                     F_Het = double(), p_Het = double())
promcomp_results_paov <- data.frame(R2_age = double(), p_colfam = double(),
                                    R2_residual = double(),
                                    F_Het = double(), p_Het = double())
highcomp_results_paov <- data.frame(R2_age = double(), p_colfam = double(),
                                    R2_residual = double(),
                                    F_Het = double(), p_Het = double())
simper_results_paov <- data.frame(R2_age = double(), p_colfam = double(),
                                  R2_residual = double(),
                                  F_Het = double(), p_Het = double())

# create list to store created objects in an iteration
iter_object_container <- list()

for (i in 1:btrap.iterations) {


  # create data.frame subsets (colony subset) by indexing the meta data.frame
  scent.f.ssb.m <- scent_factors[scent_factors$BeachAge == "SSB_1",]
  scent.f.fwb.m <- scent_factors[scent_factors$BeachAge == "FWB_1",]
  scent.f.ssb.p <- scent_factors[scent_factors$BeachAge == "SSB_2",]
  scent.f.fwb.p <- scent_factors[scent_factors$BeachAge == "FWB_2",]

  # int vector of row index number of permuted scent.ssb data.frame
  # row numbers will be used to create a permuted data.frame of
  # evenly distributed draws of individuals
  permute_rows_ssb_m <- sample(scent.f.ssb.m$SampleIndex, df.permutations, replace = T)
  permute_rows_fwb_m <- sample(scent.f.fwb.m$SampleIndex, df.permutations, replace = T)
  permute_rows_ssb_p <- sample(scent.f.ssb.p$SampleIndex, df.permutations, replace = T)
  permute_rows_fwb_p <- sample(scent.f.fwb.p$SampleIndex, df.permutations, replace = T)

  # create overall index number that can be used to
  #index data.frame(scent): index corresponds to correct individual
  perm_index_all <- c(permute_rows_ssb_m,
                      permute_rows_fwb_m,
                      permute_rows_ssb_p,
                      permute_rows_fwb_p)
```

```r
# create new data.frame with indeces found in permutation
# results vector perm_index_all
scent.permute <- scent[perm_index_all,]
scent_factors.permute <- scent_factors[perm_index_all,]
# rownames(scent.permute) == rownames(scent_factors.permute) # TRUE
#----------------------------------------

# Perform analysis to find 3 subsets based on different premises
# with the permuted data frame.
# Track 15 best performing compounds of an analysis
#----------------------------------------

## NDMS scale results
## count number of peaks that are not 0 per column
peak_count <- as.vector(apply(scent.permute, 2, function(x) length(x[x>0])))

## add peaks in a column that are not 0 to estimate highest
# concentration peak sum
peak_add <- as.vector(apply(scent.permute, 2, function(x) sum(x)))

## create dataframe with same name properties as scent.RData
compound_subset <- data.frame(name = colnames(scent.permute),
                              peak_count, peak_add)

## sort data frame for most prominent compounds over all samples
most_abundant <- compound_subset %>% arrange(desc(peak_count))

## shorten scent matrix to only the 15 most abundant compounds
scent.promcomp <- scent.permute[colnames(scent.permute) %in%
                                  most_abundant$name[1:15]]

## sort data frame for most highly concentrated compounds over all samples
most_concentration <- compound_subset %>% arrange(desc(peak_add))

## shorten scent matrix to only the 15 most abundant compounds
scent.highcomp <- scent.permute[colnames(scent.permute) %in%
                                  most_concentration$name[1:15]]

## simper
# simper analysis and results array
sim <- with(scent_factors.permute,
            simper(scent.permute, colony))
best.compounds.simper.btrap <- summary(sim)[[1]]
#filter 15 compounds that contribute most towards dissimilarity of individuals
simper_comps <- as.numeric(rownames(best.compounds.simper.btrap))
best_comps <- simper_comps[1:15]
# subset peak data matrix {scent}
scent.simper.btrap <- scent.permute[,which(colnames(scent.permute) %in%
                                             as.character(best_comps))]
#----------------------------------------

# Take 15 identified compounds and limit nMDS of the permuted
# data frame (scent.permute) to only those compounds
```

```r
#----------------------------------------

# bray-curtis similarity
scent_nmds_regular.obj <- vegan::metaMDS(comm = scent.permute, k = 2,
                                         try = df.permutations, distance = "bray")
scent_nmds_count.obj <- vegan::metaMDS(comm = scent.promcomp, k = 2,
                                       try = df.permutations, distance = "bray")
scent_nmds_add.obj <- vegan::metaMDS(comm = scent.highcomp, k = 2,
                                     try = df.permutations, distance = "bray")
scent_nmds_simper.obj <- vegan::metaMDS(comm = scent.simper.btrap, k = 2,
                                        try = df.permutations, distance = "bray")

## get x and y coordinates
scent_nmds_regular <- as.data.frame(scent_nmds_regular.obj[["points"]])
scent_nmds_count <- as.data.frame(scent_nmds_count.obj[["points"]])
scent_nmds_add <- as.data.frame(scent_nmds_add.obj[["points"]])
scent_nmds_simper <- as.data.frame(scent_nmds_simper.obj[["points"]])

## add the colony as a factor to each sample
scent_nmds <- data.frame(MDS1r = scent_nmds_regular[["MDS1"]],
                         MDS2r = scent_nmds_regular[["MDS2"]],
                         MDS1c = scent_nmds_count[["MDS1"]],
                         MDS2c = scent_nmds_count[["MDS2"]],
                         MDS1a = scent_nmds_add[["MDS1"]],
                         MDS2a = scent_nmds_add[["MDS2"]],
                         MDS1s = scent_nmds_simper[["MDS1"]],
                         MDS2s = scent_nmds_simper[["MDS2"]],
                         age = scent_factors.permute[["age"]],
                         colony = scent_factors.permute[["colony"]],
                         family = scent_factors.permute[["family"]],
                         BeachAge = scent_factors.permute[["BeachAge"]]
)
#----------------------------------------

# Perform PERMANOVA on distance matrix based limited scent compounds data
#----------------------------------------

# not subsetted
nonsubset.df_permanova <- adonis(scent.permute ~ age + colony + colony:family,
                                 data = scent_factors.permute,
                                 permutations = 9999)
nonsubset.df_hetgeneity <- anova(betadisper(vegdist(scent.permute),
                                            scent_factors.permute$colony))

# track important values of statistical analysis in this run
nonsubset_iter_res_paov <- cbind(R2_age = nonsubset.df_permanova$aov.tab$R2[1],
                                 R2_colony = nonsubset.df_permanova$aov.tab$R2[2],
                                 R2_famcol = nonsubset.df_permanova$aov.tab$R2[3],
                                 R2_residual = nonsubset.df_permanova$aov.tab$R2[4],
                                 F_Het = nonsubset.df_hetgeneity$`F value`[1],
                                 p_Het = nonsubset.df_hetgeneity$`Pr(>F)`[1])

# bind run values to track changes over iterations in the for-loop
```

```
nonsubset_results_paov <- rbind(nonsubset_results_paov,
                                nonsubset_iter_res_paov)

#prom comps
promcomp.df_permanova <- adonis(scent.promcomp ~ age + colony + colony:family,
                               data = scent_factors.permute,
                               permutations = 9999)
promcomp.df_hetgeneity <- anova(betadisper(vegdist(scent.promcomp),
                                           scent_factors.permute$colony))

promcomp_iter_res_paov <- cbind(R2_age = promcomp.df_permanova$aov.tab$R2[1],
                               R2_colony = promcomp.df_permanova$aov.tab$R2[2],
                               R2_famcol = promcomp.df_permanova$aov.tab$R2[3],
                               R2_residual = promcomp.df_permanova$aov.tab$R2[4],
                               F_Het = promcomp.df_hetgeneity$`F value`[1],
                               p_Het = promcomp.df_hetgeneity$`Pr(>F)`[1])

promcomp_results_paov <- rbind(promcomp_results_paov,
                              promcomp_iter_res_paov)

# high comps
highcomp.df_permanova <- adonis(scent.highcomp ~ age + colony + colony:family,
                               data = scent_factors.permute,
                               permutations = 9999)
highcomp.df_hetgeneity <- anova(betadisper(vegdist(scent.highcomp), scent_factors.permute$colony))

highcomp_iter_res_paov <- cbind(R2_age = highcomp.df_permanova$aov.tab$R2[1],
                               R2_colony = highcomp.df_permanova$aov.tab$R2[2],
                               R2_famcol = highcomp.df_permanova$aov.tab$R2[3],
                               R2_residual = highcomp.df_permanova$aov.tab$R2[4],
                               F_Het = highcomp.df_hetgeneity$`F value`[1],
                               p_Het = highcomp.df_hetgeneity$`Pr(>F)`[1])

highcomp_results_paov <- rbind(highcomp_results_paov,
                              highcomp_iter_res_paov)
# SIMPER
simper.df_permanova <- adonis(scent.simper.btrap ~ age + colony + colony:family,
                             data = scent_factors.permute,
                             permutations = 9999)
simper.df_hetgeneity <- anova(betadisper(vegdist(scent.simper.btrap), scent_factors.permute$colony))

simper_iter_res_paov <- cbind(R2_age = simper.df_permanova$aov.tab$R2[1],
                             R2_colony = simper.df_permanova$aov.tab$R2[2],
                             R2_famcol = simper.df_permanova$aov.tab$R2[3],
                             R2_residual = simper.df_permanova$aov.tab$R2[4],
                             F_Het = simper.df_hetgeneity$`F value`[1],
                             p_Het = simper.df_hetgeneity$`Pr(>F)`[1])

simper_results_paov <- rbind(simper_results_paov,
                            simper_iter_res_paov)
#----------------------------------------
```

```r
    # # pack all this in a list to be later on stored in a list that can be saved again
    # create name giving the iteration step
    iteration_count <- paste0("iter_", i)

    # create list that stores relevant workspace elements for an iteration step
    iter_objects <- list(scent.permute = scent.permute,
                         scent_factors.permute = scent_factors.permute,
                         scent.promcomp = scent.promcomp,
                         scent.highcomp = scent.highcomp,
                         sim = sim,
                         scent.simper.btrap = scent.simper.btrap,
                         scent_nmds_regular.obj = scent_nmds_regular.obj,
                         scent_nmds_count.obj = scent_nmds_count.obj,
                         scent_nmds_add.obj = scent_nmds_add.obj,
                         scent_nmds_simper.obj = scent_nmds_simper.obj,
                         scent_nmds_regular = scent_nmds_regular,
                         scent_nmds_count = scent_nmds_count,
                         scent_nmds_add = scent_nmds_add,
                         scent_nmds_simper = scent_nmds_simper,
                         promcomp.df_permanova = promcomp.df_permanova,
                         promcomp.df_hetgeneity = promcomp.df_hetgeneity,
                         highcomp.df_permanova = highcomp.df_permanova,
                         highcomp.df_hetgeneity = highcomp.df_permanova,
                         simper.df_permanova = simper.df_permanova,
                         simper.df_hetgeneity = simper.df_hetgeneity)

    # save everything as a list in a container list, that stores
    # information/elements of all iteration steps
    iter_object_container[[i]] <- iter_objects
    names(iter_object_container)[i] <- iteration_count

  } # end i

  paov_r2_results <- list(regular = nonsubset_results_paov,
                          promcomp = promcomp_results_paov,
                          highcomp = highcomp_results_paov,
                          simper_res = simper_results_paov)
  return(list(paov_r2_results = paov_r2_results,
              iter_object_container = iter_object_container))
} # end function
```

### Session information

```
## R version 3.5.3 (2019-03-11)
## Platform: x86_64-w64-mingw32/x64 (64-bit)
## Running under: Windows 10 x64 (build 18362)
##
## Matrix products: default
##
## locale:
## [1] LC_COLLATE=German_Germany.1252  LC_CTYPE=German_Germany.1252
## [3] LC_MONETARY=German_Germany.1252 LC_NUMERIC=C
## [5] LC_TIME=German_Germany.1252
```

```
##
## attached base packages:
## [1] stats     graphics  grDevices utils     datasets  methods   base
##
## other attached packages:
##  [1] pairwiseAdonis_0.0.1 cluster_2.0.7-1      forcats_0.5.0
##  [4] stringr_1.4.0        dplyr_0.8.5          purrr_0.3.3
##  [7] tidyr_1.0.2          tibble_3.0.0         tidyverse_1.3.0
## [10] ggbeeswarm_0.6.0     ggplot2_3.3.0        readr_1.3.1
## [13] vegan_2.5-6          lattice_0.20-38      permute_0.9-5
## [16] GCalignR_1.0.2
##
## loaded via a namespace (and not attached):
##  [1] Rcpp_1.0.4.6     lubridate_1.7.4  assertthat_0.2.0 digest_0.6.25
##  [5] cellranger_1.1.0 R6_2.3.0         backports_1.1.2  reprex_0.3.0
##  [9] evaluate_0.12    httr_1.4.1       pillar_1.4.3     rlang_0.4.5
## [13] readxl_1.3.1     rstudioapi_0.11  Matrix_1.2-15    rmarkdown_1.11
## [17] labeling_0.3     splines_3.5.3    munsell_0.5.0    broom_0.5.5
## [21] compiler_3.5.3   vipor_0.4.5      modelr_0.1.6     xfun_0.4
## [25] pkgconfig_2.0.2  mgcv_1.8-27      htmltools_0.3.6  tidyselect_0.2.5
## [29] fansi_0.4.0      crayon_1.3.4     dbplyr_1.4.3     withr_2.1.2
## [33] MASS_7.3-51.1    grid_3.5.3       nlme_3.1-137     jsonlite_1.6.1
## [37] gtable_0.2.0     lifecycle_0.2.0  DBI_1.0.0        magrittr_1.5
## [41] scales_1.0.0     cli_2.0.2        stringi_1.2.4    fs_1.4.1
## [45] xml2_1.3.1       ellipsis_0.3.0   generics_0.0.2   vctrs_0.2.4
## [49] tools_3.5.3      glue_1.4.0       beeswarm_0.2.3   hms_0.5.3
## [53] parallel_3.5.3   yaml_2.2.0       colorspace_1.3-2 rvest_0.3.5
## [57] knitr_1.21       haven_2.2.0
```
