## Supplementary figure 1 for "Chemical patterns of colony membership and mother-offspring similarity in Antarctic fur seals are reproducible over time"

**Supplementary figure 1.** Two-dimensional non-metric multidimensional scaling (NMDS) plot of chemical data from skin swabs of Antarctic fur seal pups from six colonies around Bird Island, South Georgia. NDMS was performed using Bray-Curtis similarity values calculated from log(x+1) transformed chemical abundance data. The scales of the two axes are arbitrary and the closer two points appear in the plot, the more similar they are chemically. Individual data points are colour-coded by colony.

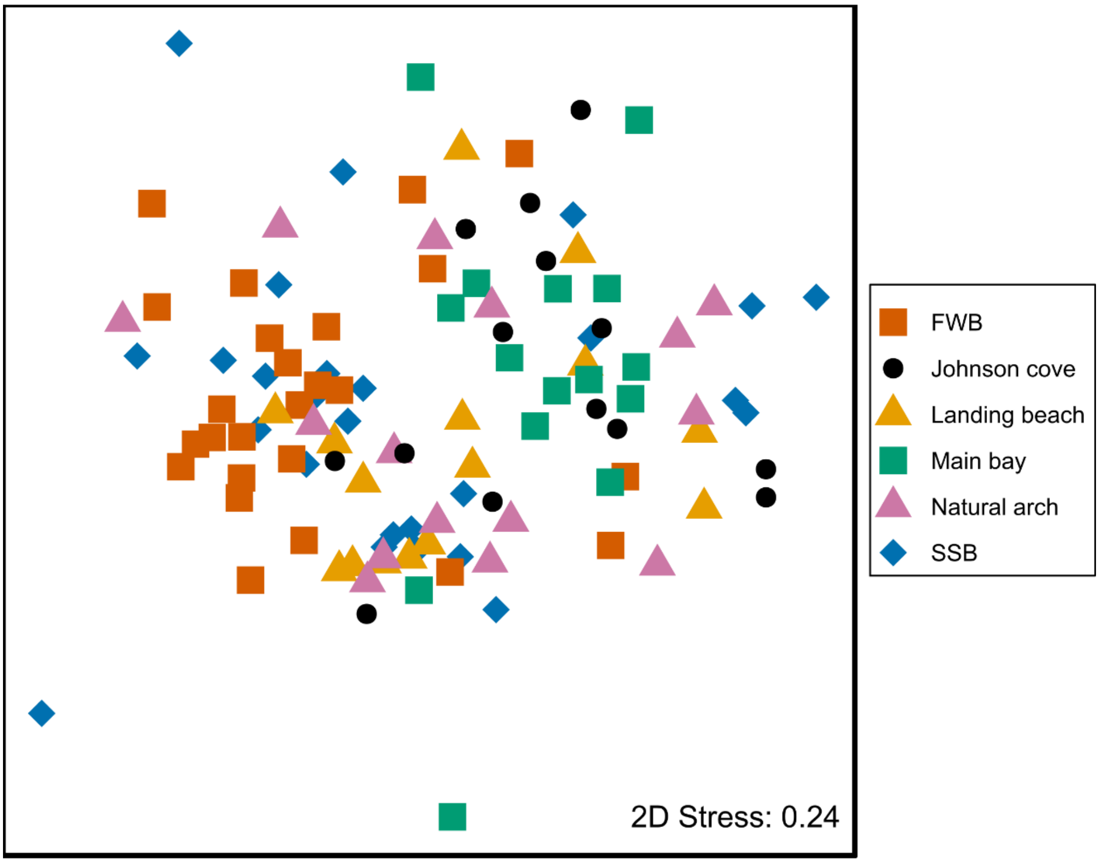
